# A shallow-scale phylogenomics approach reveals parallel patterns of diversification among sympatric populations of cryptic Neotropical aquatic beetles (Coleoptera: Noteridae)

**DOI:** 10.1101/2023.09.28.559972

**Authors:** S.M. Baca, G.T. Gustafson, D.A. DeRaad, A. Alexander, P.M. Hime, A.E.Z. Short

## Abstract

The *Notomicrus traili* species group (Coleoptera: Noteridae) is a lineage of aquatic beetles distributed throughout South America and extends into Mexico and the West Indies. Previous research has revealed a species complex within this group, with multiple distinct clades sharing overlapping distributions and lineages attributed to *N. traili* and the closely related *N. gracilipes* recovered as polyphyletic. Here, we perform targeted capture of ultraconserved elements (UCEs) to examine relationships and patterns of evolution within the *N. traili* group. First, we use short-read whole genome sequencing of four noterid genera to design a noterid-specific UCE probe set (Noteridae 3.4Kv1) targeting over 3,400 unique loci. Using this probe set, we capture UCE data from population-level sampling of 44 *traili* group specimens from across the Neotropics, with an emphasis on the Guiana Shield where distributions of several putative *N. traili* group populations overlap. We subject the resulting data matrix to various trimming and data completeness treatments and reconstruct the phylogeny with both concatenated maximum likelihood and coalescent congruent methods. We recover robust phylogenetic estimates that identify several phylogenetically distinct clades within the *traili* group that share overlapping distributions. To test for the genetic distinctiveness of populations, we extract single nucleotide polymorphism (SNP) data from UCE alignments and examine patterns of genetic clustering using principal component analyses (PCAs) and STRUCTURE. Population genetic results are highly concordant with recovered phylogenetic structure, revealing a high degree of co-ancestry shared within identified clades, contrasting with limited ancestry sharing between clades. We recover a pattern consistent with repeated diversification and dispersal of the *traili* group in the Neotropics, highlighting the efficacy of a tailored UCE approach for facilitating shallow-scale phylogenetic reconstructions and population genetic analyses, which can reveal novel aspects of coleopteran phylogeography.

## Introduction

The relationship between geography and patterns of organismal diversity has long been considered in the study of Neotropical biota (Rull, 2020), with investigations varying in focal spatial and temporal scales (e.g. Harvey et al., 2016; 2017; Toussaint and Gillet, 2018; Maestri and Duarte, 2020; Rull, 2020; Sánchez-Herrera et al., 2020; Johnson; et al., 2023). However, biogeographic hypotheses explaining the high levels of Neotropical biodiversity often focus on recent timescales (e.g. Haffer 1969; Marshall et al., 1982; Colinvaux et al. 2000; see Antonelli, 2018 and Rull, 2020). Indeed, phylogeography and reconstructions of shallow-scale evolutionary history provide valuable insights into the study of biodiversity and the dynamics of diversification, including contributions from taxonomy and species delimitation (e.g. Branstetter and Longino, 2019; 2022; Gueuning et al., 2020; Sánchez-Herrera et al., 2020). Despite comprising the greatest diversity of any group of organisms, insects are the subject of relatively few biogeographic studies within the neotropics outside of Lepidoptera (e.g. Garzón-Orduña et al., 2014; Chazot et al., 2018; Brower and Garzón-Orduña, 2020) or Hymenoptera (e.g. Branstetter & Longino, 2019; 2022; Barrera et al, 2022). This is in large part due to substantial gaps in taxonomic documentation or inadequate sampling resolution (Basset et al., 2017; Antonelli et al., 2018). Recently, aquatic beetles have seen increased utility as models in evolutionary biology (Short, 2018; Bilton et al., 2019), including in studies of biogeography (e.g. Bukontaite, et al., 2014; Toussaint and Short, 2016; Gustafson and Miller, 2017; Baca et al, 2020; Short et al., 2021; see Bilton et al., 2019 and Short, 2019), due to an increasingly comprehensive documentation of the overall diversity of the group and ongoing advances in phylogenetic reconstruction. However, most of these works incorporating Neotropical bio- or phylogeography focus on relatively deeper timescales (>20mya), with investigations of more recent phylogeography limited to the western Palearctic (e.g. Hidalgo-Galiana and Ribera, 2011; GarcíaLVázquez et al., 2016; 2017). Thus, there is a need for a more complete understanding of the recent diversification history of Neotropical aquatic beetles. This foundational evolutionary context will be key for establishing aquatic beetles as model system for testing hypotheses of Neotropical phylogeography and contributing to our understanding of Neotropical diversification. Here we combine phylogenetic and population genetic approaches in an effort to reconstruct the recent evolutionary history of the *Notomicrus traili* species group, an emerging model system within this group of Neotropical aquatic beetles.

*Notomicrus* Sharp is a genus of minute, aquatic beetles in the family Noteridae (Coleoptera: Adephaga). The genus is distributed across the New World, Oceania and Indomalaya, although most *Notomicrus* diversity is found in the Neotropics. Recent investigations have described several new species (Manuel, 2015; Baca and Short, 2018; 2021; Guimarães and Ferreira-Jr, 2019) with several more yet undescribed (Baca and Short, 2020; 2021). A species-level phylogenetic reconstruction (Baca and Short, 2020) and taxonomic circumscription of *Notomicrus* (Baca and Short, 2021) show that the genus contains several species groups, most requiring taxonomic revision. The *Notomicrus traili* species group (hereafter “*traili* group”; sensu Baca & Short, 2020:8) is Neotropical in distribution, ranging from Mexico to Argentina and into the Antilles. Baca and Short (2020) noted a high degree of phylogenetic structure within the group, including many distinct, but morphologically cryptic lineages with overlapping distributions. They also showed that characters used for morphological delimitation of species in this group (e.g. Young, 1978; Manuel, 2015; Baca & Short, 2018) were not consistent within recovered clades, i.e., the *traili* group consisted of a species complex with inconsistent geographic patterning. Additionally, individuals morphologically attributable to *N. traili* Sharp 1882 and *N. gracilipes* were recovered as polyphyletic suggesting the need for a comprehensive genomic assessment of species limits in this group.

The dynamic and complex geography of the Neotropics has been shown to strongly influence the diversification of many organisms (Hoorn & Wesselingh, 2011; Antonelli et al., 2018; Rull, 2020). While there have been several hypotheses presented to explain the abundance of Neotropical diversity (e.g. Haffer, 1969; Marshall et al., 1982), current perspectives generally acknowledge that no single generalized biogeographic hypothesis can capture the nuance of diversification across Neotropical groups (Antonelli and Sanmartín 2011; Hughes et al. 2012; Antonelli et al., 2018; Rull 2020). With apparent ecological shifts in Notomicrinae, including within the *traili* group (e.g. seep-dwelling *N. petrareptans* Baca and Short, 2018, see Baca and Short, 2020), the complex evolutionary history of *Notomicrus* is no exception. However, examining the evolutionary influence of Neotropical geography establishes essential context for understanding other aspects of *Notomicrus* diversification. Further, *Notomicrus* has previously proven effective as a system to test hypotheses related to Neotropical biogeography, albeit at deeper time-scales (Baca and Short, 2020). Given the complex history of the Neotropics with many potential geographic influences (Antonelli et al., 2018), the relationships and widespread and overlapping distributions of the *traili* group present a potential model system for developing a greater understanding of the role of geography in shallow-scale diversification in the Neotropics.

Though not a ‘cure-all’ for the challenges of elucidating patterns at different time scales, the advent of genomics has given investigators access to unprecedented amounts of data and a wealth of bioinformatic methods for untangling evolutionary histories. The targeted capture of Ultraconserved Elements (UCEs; Faircloth et al., 2012) exemplifies the increasing accessibility of genome-scale data for non-model taxa. UCEs provide systematists with a locus type that is broadly applicable to evolutionary questions across a wide breadth of evolutionary scales, from deep (e.g. Faircloth et al., 2013; Branstetter et al., 2017; Gustafson et al., 2020) to shallow (e.g. Smith et al., 2014; Harvey et al., 2016; Manthey et al., 2016; Branstetter & Longino, 2019; 2021; Gueuning et al., 2020), and incorporates a well-established, yet augmentable, phylogenomic workflow. Recent investigations have in particular shown the utility of UCEs for species delimitation and phylogeography in insects and other arthropods (e.g. Branstetter and Longino, 2019; 2021; Gueuning et al., 2020; Newton et al., 2023). Though UCEs present an ideal tool for reconstructing the evolutionary history of insects, the mechanisms of molecular evolution among and within populations and diversifying lineages are complex (e.g., incomplete lineage sorting (ILS) and gene flow; Maddison, 1997; Edwards and Beerli, 2000; Kabutko and Degnan, 2007; Edwards, 2009; Sukumaran and Knowles, 2017). While these mechanisms may influence the evolutionary history of organisms at all timescales (Oliver., 2013; Hime et al., 2021), parsing patterns and testing for processes of diversification of closely related lineages often requires a nuanced blend of population genomics and phylogenomics that runs the gamut from assignment of ancestry proportions in individual samples to large scale reconstructions of the evolutionary history of the recovered lineages (e.g., Derkarabetian et al. 2019; Derkarabetian et al. 2022; DeRaad et al. 2022). As the *N. traili* species group presents a case of hierarchical population genetic structure that blurs the lines between species and populations, it is desirable to examine patterns of evolution with an added population genetic approach, as has been successfully done in other systems using UCEs (E.g. McCormack et al., 2016; Newton et al., 2023).

Here, we use the targeted capture of UCEs to investigate evolutionary patterns among populations in the *traili group*. First, we use whole genome sequencing of select noterid taxa to design a custom, noterid-specific UCE probe set. We use this probe set to capture UCEs from individuals across the *traili* group, targeting all described species with broad geographic sampling, emphasizing the Guiana Shield and surrounding areas where much of the range overlap in the group occurs. We then reconstruct the phylogeny of the species complex, testing the effect of different combinations of data matrix trimming, completeness, partitioning, and inference methods. To further examine geographic structure within the complex, we harvest single nucleotide polymorphisms (SNPs) from our UCE alignments to use as input for genetic clustering analyses. To call SNPs from UCE alignments we extended previous approaches (McCormack et al. 2016; DeRaad et al. 2023), adding the novel step of constructing a chimeric UCE reference sequence set so that all captured UCEs across all sampled taxa are available for read mapping and SNP calling. This method effectively maximized the size of our SNP dataset, despite the lack of availability of an appropriate reference genome for this group. This is conceptually similar to the approach of Hird et al. (2011) for producing a ‘provisional reference genome’. Finally, we applied genetic clustering methods to the resulting SNP datasets to assign genomic ancestry to individual samples without the need for *a priori* sample classifications. These methods sum to an approach that (1) robustly examines the distinctiveness of lineages within the *traili group*, thereby testing the validity of described species and informing our understanding of the breadth of diversity within the complex; (2) retests the apparent polyphyly associated with external morphological characters used to delimit species in the complex; (3) allows us to examine relationships and patterns of genetic ancestry as they relate to neotropical geography; and (4) tests the efficacy of our tailored UCE probe set in application to population genetic approaches designed to identify geographic substructuring among intraspecific populations. These objectives collectively construct a key scaffold for taxonomic revision within the complex, provide a foundation for rigorous testing of the mechanisms driving evolution in *Notomicrus* and other Neotropical systems.

## Material and methods

### UCE probe set design

#### Sampling

To design a noterid-specific probe set, we selected four species for whole genome sequencing, each representing one of four genera spanning disparate phylogenetic positions within the family (Baca et al., 2017b): *Neohydrocoptus, Liocanthydrus*, *Suphisellus*, and *Sternocanthus* (Fig. 1). DNA was extracted from fresh specimens stored at -20° C in 95% ethanol with a Qiagen DNeasy kit (Qiagen, Hilden, Germany) following manufacturer’s protocols modified by the recommendations of Cruaud et al., (2018) to maximize DNA yield. Extracted DNA was quantified on a Quantus flourometer (Promega, Maddison, WI, USA) using QuantiFlour dsDNA reagents. 500ng of DNA was dehydrated and sent to RAPiD Genomics (Gainesville, FL, USA) for library prep and sequencing. Library preparation was performed by RAPiD Genomics for Illumina sequencing utilizing their high-throughput workflow with proprietary chemistry. Briefly, DNA was sheared to a mean fragment length of 350 base-pair (bp), fragments were end-repaired, followed by incorporation of unique dual-indexed Illumina adaptors and PCR enrichment. Samples were pooled at equimolar concentrations and sequenced on an Illumina HiSeq X to generate paired-end 150bp reads.

**Figure 1.**
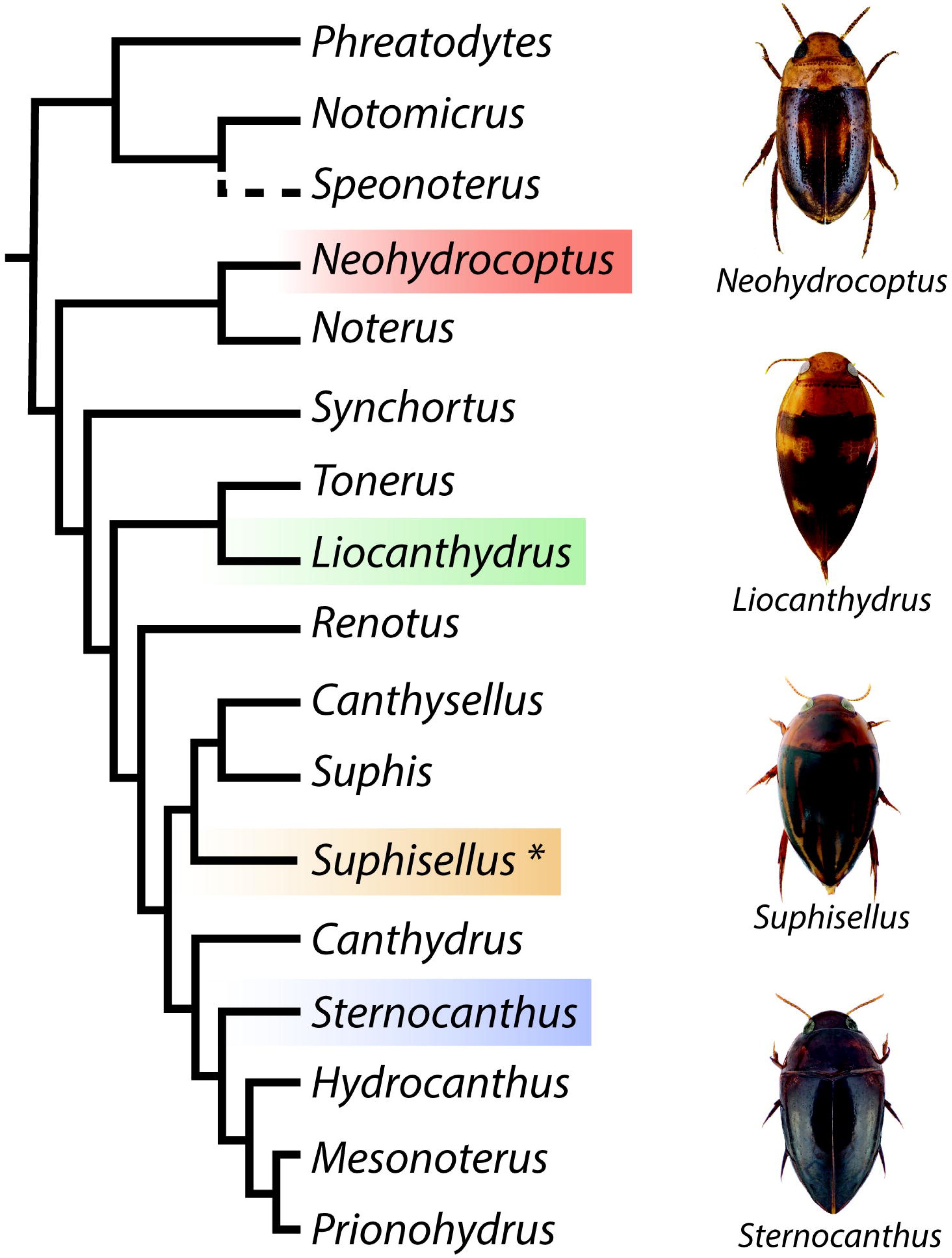
Phylogenetic positions of genera selected (highlighted) for short-read whole genome sequencing for tailored design of the Noteridae 3.4Kv1 probeset. Summarized phylogeny from Baca et al., 2017b). Image credits: S. Baca; U. Schmidt (cc by 2.0; see acknowledgements for details). Asterisk (*) indicates genus used in *sensu lato*.

#### Data processing and assembly

Demultiplexed reads for each sample were iteratively trimmed for adapters and contamination with Trimmomatic (Lohse et al., 2012) via IlluminaProcessor (Faircloth, 2013) and FastP (0.14.1 (Chen et al., 2018) with default settings. Trimmed reads were inspected for quality and adapter contamination in the FastP outputs and with FastQC (Andrews, 2010), then assembled into contigs and scaffolded with SPAdes 3.13.1 (Bankevich et al., 2012), with default kmer settings (k = 21 – 77) and read error correction. Assemblies were assessed by standard output metrics (e.g. scaffold size, N50, L50) calculated by QUAST (Gurevich et al., 2013) and by estimating genome completeness (%) with Benchmarking Universal Single Copy Orthologs (BUSCOs) using BUSCO 3.0 (Simão et al., 2015) with the endopterygota_odb9 ortholog database and the annotated *Tribolium castaneum* (Herbst) (Tribolium Genome Sequencing Consortium, 2008).

Probe set design was conducted with PHYLUCE 1.5 for Python 2.7 (Faircloth, 2016) following Gustafson et al., (2019). Gustafson et al. (2019) found that the average genetic distance of the ‘base genome’ from the other genomes used for probe design was negatively correlated with *in silico* data capture performance. To test for this pattern in our dataset, we calculated the pairwise genetic distances of Sanger loci from previously sequenced individuals (Baca et al., 2017b; *COI*, *CAD*, *H3*, *16S*, *18S*, *28S*) and from BUSCOs, which revealed that *Liocanthydrus* minimized the average pairwise distance when used as the base genome. The output probe set was then filtered for potential paralogs using a custom R script (R Core Team; Alexander, 2018; Gustafson 2019) and BLAST (Altschul et al., 1990). To further reduce the probe set, probes were selected at random with a custom R script (Alexander 2019, further_whittling_random.R). This resulting probe set was merged with probes from the Adephaga-specific Adephaga 2.9kv1 probe set (Gustafson et al., 2019, 2020) after being filtered to include only probes that (1) successfully captured the respective target UCE loci from all four noterid genomes *in silico* and (2) additionally did not target the same loci as the new noterid-specific probes. Finally, “legacy markers” (Branstetter et al., 2017b) used in previous studies of Noteridae (Baca et al., 2017b; *COI*, *CAD*, *H3*, *16S*, *18S*, *28S*) were added, resulting in the Noteridae 3.2kv1 probeset.

### UCE data capture and phylogenetics

#### Taxon sampling

We sampled 44 individuals within the *traili* group (Table 1), in addition to three outgroup taxa representing the three other species groups of New World *Notomicrus* (see Baca et al, 2020; 2021): *N. josiahi* (Miller, 2013); *N. nanulus* (LeConte, 1863); *N.* sp. 7 (of the *meizon* group, sensu Baca *et al*., 2020). Our sampling within the complex targeted (1) samples of individuals attributable to described species (including paratypes of *N. sabrouxi* Manuel, 2015 and *N. petrareptans* Baca and Short, 2018) or putative species (e.g. *N.* sp. 3, SLE895) and (2) a high resolution geographic/population sampling for these ‘species’. Sampling was iterative and guided by initial select Sanger screening of the COI mtDNA marker to ensure placement in *traili* group; COI sequencing followed methods of Baca *et al*. (2020). Sampling was emphasized in northern South America, particularly in the Guiana Shield, following these results.

**Table 1.** Summary statistics of genome assemblies used in probe set design. Genome size, mean coverage, N50 and L50 were estimated from genome assemblies using QUAST (Gurevich et al., 2013); BUSCO completeness was estimated with BUSCO 3.0 (Simão et al., 2015).

| Taxon | Genome size<br>(Megabases) | BUSCO<br>completeness | Mean coverage<br>(depth) | N50<br>(basepairs) | L50<br>(basepairs) |
| --- | --- | --- | --- | --- | --- |
| *Neohydrocoptus* sp. | 350.8 | C:79.3%<br>[S:78.4%,D:0.9%] | 8.7X | 7162 | 8720 |
| *Liocanthyrus*<br>*bicolor* | 297.7 | C:85.8%<br>[S:85.0%,D:0.8%] | 9.2X | 14047 | 4428 |
| *Suphisellus*<br>*puncticollis* | 383.9 | C:69.3%<br>[S:68.6%,D:0.7%] | 7.5X | 4518 | 23916 |
| *Sternocanthus* sp. | 578.7 | C:67.0%<br>[S:66.3%,D:0.7%] | 8.2X | 5178 | 22878 |

#### UCE data capture

Samples were extracted and quantified in the same manner as described above for samples used in UCE probe set design. Between 10ng and 120ng of DNA from each sample was dehydrated and sent to RAPiD Genomics for Library prep and enrichment with the tailored Noteridae 3.2kv1 UCE probeset. The probeset was synthesized at 2X coverage (i.e. two copies of each probe) to attempt to maximize depth of sequencing coverage for UCE loci. Library preparation was performed by RAPiD Genomics for Illumina sequencing utilizing their high-throughput workflow with proprietary chemistry. Briefly, DNA is sheared to a mean fragment length of 350bp, fragments are end-repaired, followed by incorporation of unique dual-indexed Illumina adaptors and PCR enrichment. Sequence capture was then performed using RAPiD Genomics proprietary chemistry and workflows. In general terms, the sequence capture process involves hybridizing a whole genome library from each sample to 120bp probes, capturing probe/DNA hybrids on streptavidin beads, washing away extraneous DNA, and performing PCR amplification on the remaining library that successfully hybridized with our custom probe set. Samples were then pooled to equimolar concentration and sequenced using 150 bp paired end sequencing on an Illumina HiSeq X

#### UCE data processing

UCE data were processed using PHYLUCE 1.7 (Faircloth, 2016) primarily following the workflows by Faircloth (https://phyluce.readthedocs.io/en/latest/tutorials/index.html) and Alexander, (https://github.com/laninsky/UCE_processing_steps). Raw reads were iteratively trimmed for adapter contamination and quality with Trimmomatic (Bolger et al., 2014) via the Illumiprocessor wrapper (Faircloth, 2013) and fastp (Chen et al., 2018). Read quality both before and after trimming was assessed via the fastp outputs and FastQC (Andrews, 2010). Reads were assembled into contigs using SPAdes (Bankevich et al., 2012) with default paired-end settings and default k values up to k = 77. Contigs were matched to UCE probes with a minimum percent identity of 80 followed by extraction of UCE loci. UCEs were aligned in PHYLUCE with MAFFT 7.215 (Katoh & Standley, 2013) with the “-- no trim” option, then internally and edge trimmed via gblocks 0.91 (Talavera & Castresana, 2007). Gblocks removes ambiguously aligned sites and gaps and trims alignment edges according to stringency thresholds. This is a common practice used to reduce alignment error or the incorporation of saturated sites. Though generally suggested for deeper evolutionary scales (Faircloth, 2016), visual inspection of select alignments showed many gaps and potential benefit from internal trimming here. We used relaxed stringency parameters (b1 0.5, b2 0.5, b3 12, b4 7) to attempt to avoid removing phylogenetically informative sites. To test the effect of gblocks, we also skipped this internal trimming step and assembled data sets with only alignment edges trimmed (by omitting the “-- no trim” option in MAFFT alignment). UCE data matrices for both treatments were then constructed at 70% and 90% taxon completeness (i.e. from UCE locus alignments with 70% and 90% representation of total taxon sampling). Missing data statistics for matrices were calculated with AMAS (Borowiec, 2016).

#### Phylogenetic analyses

For each of the four combinations of data trimming and taxon completeness treatments we performed, we output a concatenated UCE alignment. To account for heterogeneity in molecular evolution both among and within UCE loci, all concatenated alignments were partitioned with the sliding-window site characteristics based on entropy (SWSC-EN) method (Tagliacollo & Lanfear, 2018), developed for UCE data. This was accomplished by exporting respective concatenated matrices with phyluce_align_format_nexus_files_for_raxml (with charset flag) for use with the SWSC-EN script which divides each UCE locus into three data subsets: the generally more conserved core and two more rapidly evolving flanks. Maximum likelihood analyses on all concatenated datasets were conducted in IQ-TREE 2.1.4 (Minh et al., 2020b). ModelFinder (Kalyaanamoorthy et al., 2017; implemented in IQ-TREE) was used to search for best fit model of evolution for all Maximum likelihood analyses. The respective best fit models for unpartitioned analyses were searched with the “-m MFP” command, while the best fit partitioning scheme and models for partitioned analyses were searched using the “-m MFP+MERGE” command, which combines similarly evolving data under a respective shared model of evolution. Branch support for phylogenetic inference was assessed with 1,000 replicates of ultrafast bootstrap approximation (UFboot; Hoang et al., 2018), with a support value ≥ 95 considered strong support for a given branch (Hoang et al., 2018).

Coalescent-congruent analyses were conducted with ASTRAL III 5.6.1 (Zhang et al., 2018), wherein a single species tree is constructed from input gene trees. To avoid phylogenetic error due to the sensitivity of coalescent-based analyses to missing data (Hosner et al., 2016), only the 90% complete matrices of both trimming treatments (edgetrimmed, and gblocks) were used as input to create gene trees using IQ-TREE. Best-fit models of site evolution for each locus were searched in ModelFinder (Kalyaanamoorthy et al., 2017) before subsequent tree inference to account for among-locus evolutionary heterogeneity. Branch support for the ASTRAL species tree was assessed by local posterior probability (Sayyari & Mirarab, 2016).

### Population genetics

To examine the distinctiveness of *traili* group lineages with overlapping distributions and thereby test the efficacy of our tailored UCE probe set for applications in population genetic methods, we used SNP data as input for multiple genetic clustering approaches. SNPs were called among the 38 samples comprising the core *traili group* (i.e., with outgroups, *N. gracilipes*, and *N.* sp. 3 samples removed). Because of the lack of an appropriate reference genome, we generated a pseudo-reference genome (e.g., McCormack et al. 2016; DeRaad et al. 2023) for read mapping and SNP calling. In brief, this approach involves the *de novo* assembly of UCE loci using phyluce (Faircloth, 2016), which can then be used as reference sequence for mapping cleaned UCE reads from each sample and performing SNP calling. To maximize SNP extraction from the UCE dataset, we extended this approach by generating a ‘chimeric reference’ sequence assembled from multiple samples to represent all unique UCE loci present across the entire dataset.

#### Chimeric reference assembly

UCEs were identified and extracted to an incomplete matrix from the assembled contigs as above (‘*UCE data processing*’). The resulting monolith fasta file was exploded to sample-specific fastas (*phyluce_assembly_explode_get_fastas_file*, with the *‘-- by taxon*’ flag). The chimeric reference was then generated using custom bash scripts (available at https://github.com/StephenBaca). First, a list of all unique UCE loci present across the data matrix was generated. Samples were sorted by data capture success as a function of the number of UCE contigs as reported in the log file of the UCE identification step (*phyluce_match_contig_to_probes*). UCE loci from the list of unique loci were then searched in sample fastas, starting with the sample with the ‘best’ data capture (in this case SLE 2061). Matched UCE sequences were added to a single ‘chimeric’ fasta and removed from the list of unique loci. Remaining loci in this list (i.e. those not identified in the first sample’s fasta) were then searched in the next sample and those matched were also added to the chimeric fasta. This continued until each unique UCE locus identified across all 38 samples was represented in the chimeric reference.

#### SNP extraction and processing

The chimeric reference sequence was indexed with BWA and cleaned UCE reads mapped with BWA-MEM before being sorted with SAMtools. Duplicate reads were marked, and BAM files indexed, with Picard. Stats were generated with SAMtools (*flagstat*) for QC pass/fail.

A sequence dictionary was prepared from the chimeric reference sequence using Picard (*CreateSequenceDictionary.jar*) which was then indexed with samtools (*faidx*). GATK 3.8.1 was used to locate indel intervals (RealignerTargetCreator), realign reads (IndelRealigner), pre-call SNPs and indels (HaplotypeCaller), genotype and merge GVCFs (GenotypeGVCFs), and call SNPs and indels (SelectVariants x2: -selectType SNP, -selectType Indel). We then extracted SNPs, masked indels, performed quality filtering (VariantFiltration: -maskExtension 5, -QUAL < 20.0, -QD < 2.0) and output a variant call format (vcf) file, with GATK 3.8.1. A bash script was used to filter this vcf file, retaining SNPs with a ‘PASS’ flag, i.e. those that passed quality filtering (“cat filtered.vcf | grep ‘PASS\|^#’ > PASSonly.vcf”). A final filtering step was applied with VCFtools (Danecek et al., 2012; settings: MAF=0.1, MISS=0.8, QUAL=20, MIN_DEPTH=5). Multiple subsampled vcf files (three separate files corresponding to Clade 1, Clade 2, Clade 3+4) were generated with VCFtools with respective sample lists passed to the -- keep and --remove flags.

We read each of the resulting subsampled (Clade 1, Clade 2, Clade 3+4) and full vcf files into R using the R package vcfR (Knaus and Grunwald, 2017). We then used these vcfR objects as input for the R package SNPfiltR (DeRaad, 2022) to automate SNP data filtering, visualize the effects of different filtering treatments, run PCAs, and prepare SNP data for downstream analyses. Following the SNPfiltR recommended pipeline, genotypes were hard filtered to a minimum depth of 3 reads, genotype quality of 25, an allele balance ratio range of 0.25–0.75, and a maximum coverage cutoff of 100 reads (to remove potential paralogous SNPs). Missing data was assessed by sample and by SNP across different filtering thresholds with a 70% (0.7) completeness threshold applied per SNP. We ran genetic clustering PCAs and cross-referenced sample clustering with missing data per sample to confirm that missing data was not driving clustering patterns at the specified cutoffs. Invariant sites resulting from these filtering steps were appropriately removed before SNP datasets were used in downstream analyses, as specified in the SNPfiltR workflow. For population assignment analyses (see ‘STRUCTURE’ below), SNP datasets were filtered for singletons in SNPfiltR and then for linkage by distance thinning to one SNP per 5000bp window in VCFtools (--thin 5000).

#### STRUCTURE

To examine genetic structure among and within populations, unlinked SNP datasets were subjected to parametric genetic clustering analyses in STRUCTURE (Pritchard *et al*. 2000), which uses sites’ allele frequencies to assign individuals to a pre-set number of populations (or genetic clusters, K) based their genotypes. Additionally, the Q parameter was invoked to allow for proportional assignment of an individual’s genome to a population. The SNP dataset was iteratively divided into sampling subsets (Fig 2; Table 5): Clade 1 (13 individuals), Clade 2 (9 individuals), Clades 3 + 4 (16 individuals), as guided by an initial run of the full SNP dataset including all 38 individuals and clades identified in phylogenetic reconstructions.

**Figure 2.**
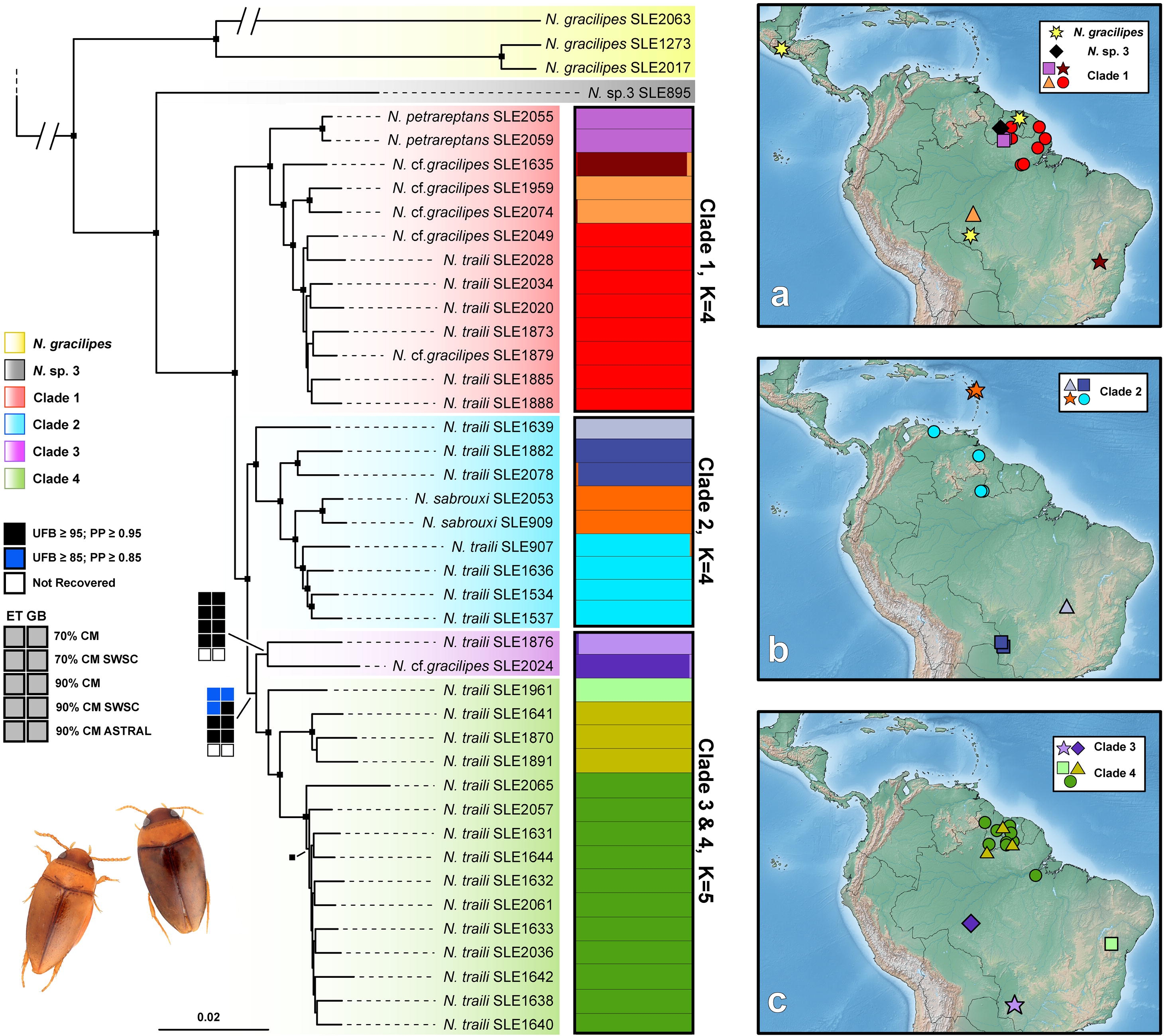
Phylogeny, admixture plots and distribution maps of the *Notomicrus traili* group. Phylogeny based on Maximum likelihood analysis of the 90% CM-SWSC matrix. Color overlay indicates respective clade as depicted in legend at left. Black boxes at nodes indicate high support for the depicted relationships across all analyses; 10-box configurations at backbone nodes depict low support or incongruence; legend at bottom left depicts analyses corresponding to boxes; color indicates support. ET = edge trim treatment; GB = gblocks treatment; SWSC = Sliding Window Site Characteristics partitioning treatment; 70% CM = 70% completeness matrix; 90% CM = 90% complete matrix. Admixture plots show population assignment of individuals in tree at optimal K values (according to the Evanno method) as recovered by STRUCTURE in analyses subsampled by clade. Maps at right show distributions of corresponding clades as legend indicates, maps ordered in descending phylogenetic order; colors of shapes in map correspond to colors in respective clades and admixture plots (e.g. in map A, *N. gracilipes* (yellow clade) corresponds to yellow stars; red population in Clade 1 admixture plot corresponds to red circles); note colors also correspond accordingly to samples in PCAs in Fig. 3. Beetles at bottom left, left to right: *N.* cf. *gracilipes*, *N.* cf. *traili*. Images: S. Baca.

STRUCTURE analyses were run across a range of K values for each prepared SNP data subset: K_Clade 1_ = 1–5, K_Clade 2_ = 1–5; K_Clade 3 + 4_ = 1–8. Analyses consisted of five independent runs across each K value for each subset, each for 500,000 reps, iteratively increased to 3 million reps for analyses of Clade 3+4 as runs converged poorly at certain K values (Francis, 2017). Performance of each run was evaluated using the pophelper (Francis, 2017) R package. Convergence of runs across K values for respective datasets was assessed via Evanno plots using the Evanno method (Evanno, 2005) as implemented in pophelper, and based on the consistency of population assignments across identical replicate runs.

## Results

### Probe design

Whole genome sequence assembly statistics for the four noterid taxa are given in Table 1, with an estimated average coverage depth of 7.5–9.2X and a BUSCO estimated completeness of 67% – 85% (Table 1). Ranked average pairwise genetic distances for Sanger loci and BUSCOs are shown in Table 2. *Liocanthydrus bicolor* was shown to have the lowest average genetic distance in both Sanger and BUSCOs data. This was concordant with the results of the base genome tests that indicated that candidate loci designed with *Liocanthydrus* as the base genome captured, *in silico*, the most loci with complete taxon representation (i.e. loci were captured across all four genomes used in probe design) (Table 2).

**Table 2.** Base genome selection for probe design. Average ranked pairwise genetic distance of each genome to others based on Sanger markers and BUSCOs (identified in BUSCO 3.0 using the endopterygota_odb9 ortholog database) and number of identified UCE loci and targeting probes in resultant probe set in base genome tests. Lowest distance values and highest loci and probe values in bold.

| Taxon | Sanger<br>markers | BUSCOs | UCE loci | Probes |
| --- | --- | --- | --- | --- |
| *Neohydrocoptus* sp. | 3.83 | 3.4 | 55,952 | 430k |
| *Liocanthyrus*<br>*bicolor* | **1.83** | **1.4** | **67,783** | **517k** |
| *Suphisellus*<br>*puncticollis* | 2.5 | 2.2 | 54,760 | 416k |
| *Sternocanthus* sp. | **1.83** | 3 | 52,812 | 400k |

The final UCE probeset was designed with *Liocanthydrus* serving as the base genome. The initial probe design yielded over 50,000 loci. After removal of potential paralogs, this was reduced to ca. 14,000 loci. With random UCE locus selection, the final Noteridae 3.4kv1 probe set targeted the following: 3,198 noterid-specific UCE loci, 11 UCE loci shared with the Adephaga 2.9kv1 probe set (Gustafson et al., 2019; 2020); 171 UCE loci from the Adephaga 2.9kv1 probe set that were captured *in silico* from all four noterid genomes, and six ‘legacy’ loci (Sanger markers). This totaled 3,385 total loci targeted by 28,000 probes; with 2x probe enrichment, 56,000 probes were used in UCE data capture.

### UCE data capture

We captured a total of 2,407 UCE loci in the incomplete matrix (excluding the ‘legacy’ markers that were being sequenced for a separate study), across all 45 taxa, with an average of 1239.91 loci per taxon, ranging from 871 to 1,344 loci per sample. Alignment of the incomplete matrix resulted in 1,689 loci, with a total length of 1,705,508 bp (after edge trimming) and 1,222,654 bp after internal trimming with gblocks. The aligned locus length averaged 1009.77 bp (edge-trimmed; 724.32 bp with gblocks) and ranged from 111bp to 7,969 bp for edge trimmed loci, and 34 bp to 1,878 bp for gblocks trimmed loci (see Table 4).

Matrix assemblies included a minimum of 31 (70%) and 40 (90%) of the 45 taxa (Table 4). The 70% complete edge-trimmed matrix (70% CM-ET) comprised 1,214 UCE loci, a total concatenated alignment length of 1,517,815 bp, and mean UCE locus length of 1009.77 bp (range: 111–7,969 bp). The 90% edge-trimmed matrix comprised 1,017 UCE loci with a total concatenated length of 1,321,125 bp. The mean length of aligned loci was 1,299.04 bp, ranging from 382–7,969 bp (Table 4). Proportions of missing data (including both gaps “-” and missing data “?”) differed relatively little among different completeness levels of the edge-trimmed matrices (36.93% and 35.73% missing data in the 70% CM-ET and 90% CM-ET, respectively; Tables 4, S1). Medians of missing data for these matrices were 34.48% and 33.07%, respectively (Tables 4, S1). As expected, the samples with the largest proportion of missing data and gaps were outgroups and dried museum samples (Table S1).

The removal of ambiguous sites, extensive gaps, and sites with poor taxon representation by gblocks reduced average alignment length significantly, even with relaxed parameters (Tables 4, S1). The 70% complete gblocks trimmed matrix (70% CM-GB) comprised 1,214 UCE loci with a total length of 1,073,799 bp. The mean length of the aligned loci was 884.17 bp, with a range of 291–1,878 bp. The 90% complete gblocks trimmed matrix (90% CM-GB) comprised 1,017 UCE loci with a total length of 933,820 bp. The mean length of the aligned loci was 918.21 bp, with a range of 380–1,878.

The gblocks-trimmed datasets showed decreases in overall missing data (gaps “-“ and missing “?”) compared to their edge-trimmed counterparts at both completeness levels, with mean missing data at 12.94% and 11.24% for the 90% CM-ET and 90% CM-GB datasets, respectively (Table 4). The gblocks matrices showed a similar trend to the edge-trimmed matrices in that levels of completeness seemed to have only a moderate effect (Tables 4, S2). The greatest effect of internal trimming was seen in the gblocks removal of gaps (“-“), with mean gaps in the 70% CMs reduced from 24.46% (median 25.80%) to 0.94% (median 0.65%) and in the 90% CMs from 24.94% (median 26.11%) to 0.95% (median 0.64%), while changes in data coded as missing “?” decreased by <1% (Tables 4, S1).

### Performance of phylogenetic analyses

All phylogenetic analyses yielded well-resolved trees with strong support and largely congruent relationships. Topological conflicts among analyses were largely restricted to the shallowest relationships within populations of densely sampled geographic areas, with most major clades and larger sub-clades/populations in the complex consistently recovered across all analyses (Figs. 2, S2–S11). There was a notable conflict between concatenated maximum-likelihood and summary species tree analyses along the backbone of the phylogeny, with respect to Clade 3. Specifically, a species tree built using ASTRAL split this clade, recovering sample SLE1876 from Mato Grosso do Sul, Brazil as sister to Clade 1 with strong support, rather than as part of Clade 3 (PP > .95; Figs. 2, S2–S11). Meanwhile, SLE2024 remained in a congruent position, sister to Clade 4. ASTRAL analyses otherwise recovered well-resolved trees with relationships congruent with concatenated analyses or only weakly supported if suggesting a conflicting topology (Figs. 2, S10, S11).

### Phylogenetics

All analyses recovered *N. gracilipes* and a putative undescribed species, *N.* sp. 3, as successive sisters to the rest of the *traili group* (Fig. 2). The remaining complex is comprised four clades recovered across all concatenated analyses, which are arbitrarily referenced as clades 1–4 (as in Fig. 2) for convenience. Among these four clades, Clade 1 is sister to the others and includes *N. petrareptans*, a seep-dwelling species endemic to Suriname, sister to a clade extending from Minas Gerais, Brazil northeast to Para, Amapá, and Suriname. Clade 2, sister to Clade 3 + 4, consists of a complex extending from the Brazilian states of Goiás and Mato Grosso do Sul, north to northern Roraima (Brazil) and Venezuela. *Notomicrus sabrouxi*, from the Antillean island of Guadeloupe, was consistently found nested in this clade, sister to the Roraima and Venezuela populations. Clade 3 comprises only two individuals from Western Brazil, Rondônia and Mato Grosso do Sul, which are together sister to Clade 4 (Fig. 2), except in ASTRAL analyses (Figs. S10, S11). Meanwhile, Clade 4 comprises several subclades/populations. A single individual from Bahia, Brazil is recovered as sister to the rest of the clade. A population from Roraima and Suriname and an individual from Para, just south of the Amazon River and near the type locality of *N. traili*, are successive sisters to a densely sampled population from the rainforests of Guyana and Suriname.

Within these main clades, strong and consistent geographic signal was seen in relationships among populations and subpopulations; with a repeating pattern of southern South American populations recovered sister to northern South American or Guiana Shield populations. Further, multiple, phylogenetically distinct Guiana Shield populations were recovered within both clades 1 and 4. In the case of clade 4, two phylogenetically distinct Guiana Shield populations were recovered as reciprocal sisters with strong support (Fig. 2).

### Population Genetics

Full stats for subsampled SNP datasets are listed in Table 5. The SNP calling and extraction workflow resulted in an output vcf file containing 30,347 SNPs across 1,622 UCE loci. After subsetting and filtering, the dataset was drastically reduced to datasets of a few thousand or less than 1,000 SNPs depending on the filtering treatment (Table 5). The thinned, unlinked datasets used in STRUCTURE analyses were all limited to one SNP per locus with singletons removed (Table 5).

Missing data was shown to have minimal effects in PCAs across by-SNP missing data thresholds (Figs. 3, S12–17), i.e., stricter filtering thresholds up to 90% completeness cutoffs showed similar trends in clustering, thus we set by-SNP thresholds to 70% to increase the number of retained SNPs in all datasets. PCAs of Clades 1 and 2 both recovered four distinct genetic clusters (Figs. 3; S12–S15); the PCA of Clade 3+4 recovered two distinct clusters in addition to three independently placed individuals (Figs. S16, S17). Recovered patterns of genetic clustering reflected population structure in phylogenetic reconstructions (Figs 3; S12– 17).

**Figure 3.**
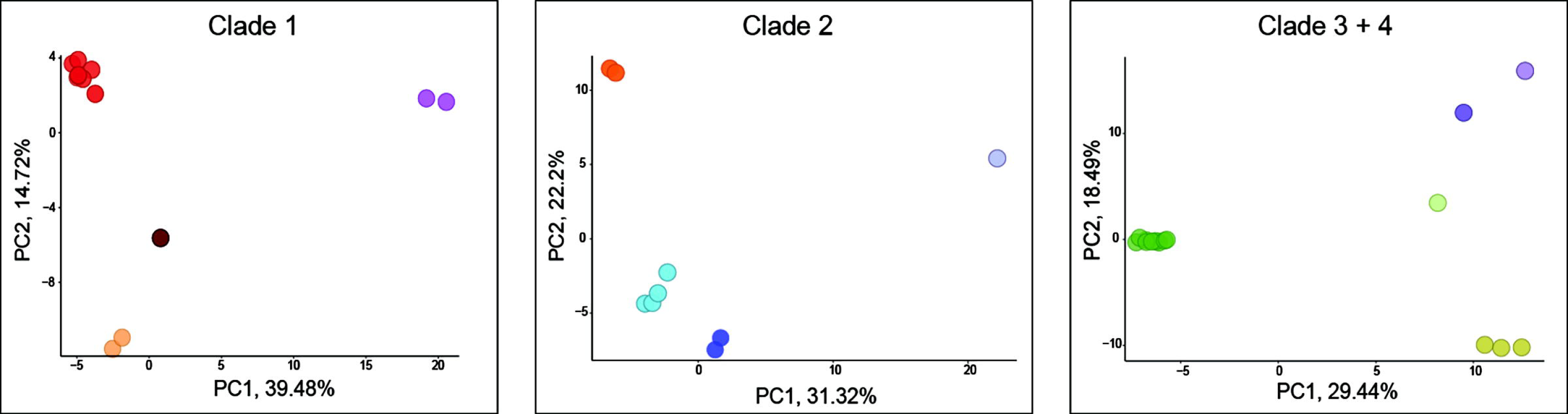
Principal component analysis (PCA) plots of SNP datasets subsampled by clade. Clades are labeled in Fig. 2; sample colors in plots correspond to population assignment in Fig. 2.

STRUCTURE analyses of Clades 1 and Clade 2 each assigned individuals to an optimal four populations (K = 4; Figs. 2, S18, S19) based on convergence across runs and support from Evanno plots (Figs. S18, S19). Analyses of Clade 3+4 with K=5 converged poorly with respect to the assignment of SLE1876, even when reps were iteratively increased to 3 million (Figs S20). Some K=5 runs assigned individuals to five distinct populations without notable admixture (as in Fig 2, while others assigned individuals to four distinct populations and SLE1876 to a fifth, admixed population, sharing variable genetic background with SLE2024 (Fig. S20). Evanno outputs showed lowered mean ln likelihood for K=5 runs, but with a very high standard deviation (Figs. S20). Inspection of STRUCTURE outputs verified high variance in ln likelihood and Ln likelihood of the data across these runs and further showed that runs with five distinct populations were recovered with the highest ln likelihood estimates, highest estimates of Ln likelihood of the data, and least variance in likelihood compared to other K=5 runs and runs at other K values. K=6 analyses also largely assigned individuals to five distinct populations, with very little admixture and almost no genetic background from a sixth population present. Overall, STRUCTURE results reflected population structure in phylogenetic reconstructions and genetic clustering in PCAs (Figs. 2, 3, S2–25).

## Discussion

We sought to clarify the relationships within the *traili* group as an emerging model system for investigations of neotropical biogeography and thereby test the efficacy of a tailored UCE probe set for shallow-scale phylogenetics and population genetics in aquatic beetle systems. We confirm that relatively low coverage (here, 7–10X) whole genome sequencing is suitable for tailored probe set design, and robust UCE data capture in insect groups with relatively few genomic resources. This is consistent with Gustafson et al., (2019; 2020; 2023); Baca et al., (2021), and Van Dam et al., (2023). Our approach proved effective in phylogenetic reconstruction at the species to population level, with phylogenetic estimates of the *traili* group recovered with high nodal support and topological congruence across data trimming and partitioning treatments and phylogenetic approaches. With these data we further extracted robust, well-curated, SNP datasets applicable to population genetic methods.

PCAs and STRUCTURE analyses recovered genetic clusters and population assignments, respectively, that reflect the relationships recovered in phylogenetic reconstructions. Together these results show that the *traili* group comprises four main clades with nested subpopulations, many of which are phylogenetically and genetically distinct despite overlapping distributions. Relationships among and within these populations show repeated geographic structuring wherein southern populations in the Brazilian Shield and Pantanal are sister to northern populations in the Guiana Shield, Antilles, and Central America. This pattern suggests parallel diversification and range expansion into subpopulations throughout the Neotropics. Importantly, we also find that the Guiana Shield harbors a high number of sympatric populations within the *traili* group. This is consistent with the historical geography of South America, in which the Guiana Shield (and Brazilian Shield) are thought to have provided relatively stable habitats compared to more dynamic areas, e.g. the Amazon Basin and Andes (Hoorn & Wesselingh, 2011 and works therein; Bicudo et al, 2019; Baker et al., 2020). Finally, our results also show that elytral punctation, a character often used for diagnosing species this group (e.g. Young, 1979; Manuel 2015; Baca and Short, 2018), shows plasticity among *traili* group lineages and should be used cautiously in species descriptions and identification. We discuss these and other aspects in detail in the following subsections.

### Performance of probe design and UCE data capture

Because UCE locus identification is based on sequence similarity, designing probes from genomes of more closely related species is expected to produce far more putative UCE loci than in other systems bridging greater scales of divergence in insects (e.g. Hymenoptera, Faircloth et al., 2015; Arachnida, Coleoptera, Diptera; Hemiptera; Lepidoptera, Faircloth, 2017; Gustafson et al., 2019; 2020, Coleoptera: Adephaga; and others, see review in Gustafson et al., 2023). Here, a high number of UCE loci were identified *in silico* during probe design (over 14K before random reduction), as expected given the assumed sequence similarity among these recently diverged species from a single family of aquatic beetles (Noteridae). It is difficult to assess potential tradeoffs associated with a high rate of UCE identification due to an increase in relative conservation. For example, it is possible that decreased sequence divergence across model genomes could increase the potential for identifying multiple UCEs from the same respective genomic locus or region, thus producing linked, congenic loci in the dataset (Van Dam et al., 2021).

The UCE data capture within *Notomicrus* species and the *traili group* using a tailored UCE probe set exceeded previous investigations of Adephaga (Baca et al., 2017; 2021; Gustafson et al, 2020) in terms of number of captured loci and data completeness, despite the absence of a *Notomicrus* model in probe design. Overall, our results are congruent with previous studies (e.g. Gustafson et al., 2019; 2020; 2023; Van Dam et al., 2023; see review in Gustafson et al., 2023) that have shown that tailored probe sets are effective for improved UCE data capture in Coleoptera. Although we are unable to construct a direct comparison for assessing the effectiveness of 2X probe tiling (i.e. synthesizing two replicates of each probe) versus 1X, the performance of data capture using our approach, along with theoretical expectations, suggests this as a viable avenue for improved UCE data capture.

### Performance of analyses

Concatenated phylogenetic reconstructions of all data completeness, trimming and partitioning regimes consistently recovered the same major clades, sub-clades and even subpopulations with largely congruent relationships, despite a large reduction in alignment sizes with internal trimming via gblocks. Given the relative conservation of UCE loci and the shallow evolutionary focus of the investigation, alignment ambiguity due to saturation would not be expected. Visual inspection of alignments of select loci confirmed that large gap-only sections were apparent. This could be caused by a combination of the relatively large genetic distance to the outgroups (e.g. *N. nanulus*) resulting in indels or non-overlapping representation in loci, and/or the effect of including dried museum specimens, and/or the default MAFFT settings in Phyluce not being optimal (see Baca et al., 2021). Trimming with gblocks appeared to largely remove and close these gaps, with almost no changes in sites coded as missing (“?”; Table S1). Alignment stats show that informative sites per locus remained constant or slightly increased after internal trimming with gblocks, suggesting little to no downside to performing this step for datasets such as ours focused on relatively shallow divergences (Table S2).

Dried museum specimens and outgroups showed the greatest levels of missing data and recovered variably long branches in phylogenetic reconstructions (Figs. 2, S2–S11, Table S2). Such effects associated with target capture studies using older museum specimens are known in insects and other arthropods (e.g. Blaimer et al., 2016; Branstetter et al., 2023; Freitas et al., 2023; Goodman et al., 2023) and are especially well documented in birds (e.g. Lim and Braun, 2016; Moyle et al., 2016; Salter et al. 2022; Hruska et al., 2023; Decicco et al., 2023; Ostrow et al., 2023). It is likely that processing captured data outside of Phyluce (e.g. mapping reads to a reference genome, aligning manually with MAFFT to adjust parameters., etc.) as in Decicco et al. (2023) could remedy some of these effects.

The difference in recovered topologies in ASTRAL analyses could be attributed to multiple potential factors. Coalescent-congruent reconstruction methods such as ASTRAL are more affected by discordance in gene trees, whether driven by poor gene tree resolution (Huang and Knowles, 2009) or gene tree estimation error (Mirarab et al., 2016), e.g. due to missing data (Hosner et al., 2016; Moyle et al., 2016), or evolutionary phenomena such as incomplete lineage sorting (ILS; Maddison, 1997; Degnan and Rosenberg, 2009) and gene flow. The short internodes among Clades 2, 3, and 4 are conducive to such effects. Note this does not indicate that the concatenated analyses should necessarily be favored over coalescent-congruent methods (see Edwards et al., 2016). In that vein, while congruence and high nodal support across trimming and partitioning regimes in our concatenated reconstructions do invoke confidence, high nodal support is generally expected in large, concatenated datasets (Liu and Edwards, 2009; Lanfear, 2018; Minh et al., 2020) even if the recovered topology is incorrect. The distribution of relationships in gene trees could also be potentially biased by congenic loci. Van Dam et al. (2021) showed that removing congenic loci increased phylogenetic performance, particularly in coalescent methods. Identifying congenic loci during the bait design phase, or potentially filtering based on locus identity (depending on the objectives of the respective investigation), or concatenating identified congenic UCE loci may be desirable. However, the methods of Van Dam et al. (2021) require an annotated reference genome which is unavailable for Noteridae or its parent suborder Adephaga at the time of this investigation (see Baca et al., 2021). It would also be desirable to screen for contamination in UCEs and the model genomes used in probe design as this has been shown to effect phylogenetic reconstruction, especially in specimens with degraded DNA, e.g. museum specimens (McCormack et al., 2016; Van Dam et al., 2021).

Robust SNP datasets were successfully extracted from UCE reads even with relatively stringent calling and filtering thresholds, including creating datasets of unlinked SNPs (filtered for one SNP per locus). Studies using similar SNP calling protocols have shown the effectiveness of SNP datasets extracted from UCE enriched reads (e.g. McCormack et al., 2016; Decicco et al., 2023; DeRaad et al., 2023; McCormack et al. 2023), including in comparisons with other reduced capture methods such as RAD-Seq (Harvey et al., 2016; Manthey et al., 2020). Studies in arthropods have commonly applied UCEs and other target capture methods for resolving deeper time-scales (E.g. Blaimer et al., 2016; Branstetter et al., 2017a, b; Starret et al., 2017; Derkarabetian et al., 2023; Homziak et al., 2023; see also Zhang et al., 2019; Gustafson et al, 2023 for more comprehensive reviews), with relatively few used in shallow scale reconstructions (e.g. Branstetter and Longino., 2019; 2022) and fewer using SNP-based analyses (Derkarabetian et al., 2019; Newton et al., 2023). A future goal of this research will be to increase the sampling within Clade 3, to better understand the genetic structure within these populations. The performance of our analyses builds shows the efficacy of combining tailored UCE probe design and reference-based SNP extraction for integrating phylogenomics and population genetics in investigations of shallow-scale evolution in other insect systems.

### Evolution of the *traili* group Relationships

We here clarified relationships within the *traili* group, finding that it comprises several cryptic lineages with sympatric distributions, and further supporting *Notomicrus* and Noteridae as models for testing biogeographic hypotheses across evolutionary time. As in Baca and Short (2020), *Notomicrus gracilipes* and *N.* sp. 3. are recovered as successive sisters to the rest of the complex. *Notomicrus gracilipes* appears to extend from northern South America and into Guatemala, where the type was collected. Further sampling is needed to assess contact and genetic structure among the highly divergent populations recovered within *N. gracilipes*. Due to phylogenetic distinctness, long branches, disjunct distributions, and scant population sampling, *N. gracilipes* and *N.* sp. 3 were omitted from downstream genetic clustering analyses.

Clade 1 relationships were largely consistent with Baca and Short (2020), especially along its backbone. *Notomicrus petrareptans* is recovered as sister to a widespread group comprising distinct populations that correlate to geography (Fig. 2, 3). Phylogenetic structure, PCAs and STRUCTURE support the distinctness of *N. petrareptans* from the rest of Clade 1, including a Guiana Shield population with an overlapping distribution. While *N. petrareptans* is genetically distinct from this sympatric population, distance and imbalanced population sampling makes difficult to assess the presence of barriers to gene flow i.e., the degree of reproductive isolation between these taxa. *N. petrareptans* is known only from a single hygropetric seep habitat in Suriname, adding an ecological correlate to its distinction. Many of the other individuals of this clade were collected from lentic and lotic habitats embedded in open, scrub and grassland areas (e.g. Sipaliwini Savannah, Surname; Amapá, Brazil; Cerrado, Minas Gerais, Brazil; Fig. 2; Table 3). Genetic clustering analyses (both PCA and STRUCTURE) concordantly recovered four distinct populations within this clade overall, which align with the internal branching pattern recovered in our phylogenetic reconstruction (Clade 1; Fig. 2, 3).

**Table 3.** Taxon sampling, including whole genome samples for UCE probe design. *Notomicrus* samples in descending order of tree tips in Fig. 2. Single asterisk (*) indicates outgroup; double asterisk (**) indicates sample was dried museum specimen.

| Taxon | Sample code | Country | State/Province | Year collected | Field code | Lat. | Long. |
| --- | --- | --- | --- | --- | --- | --- | --- |
| *Sternocanthus* sp. | SLE1544 | Madagascar | Antsiranana | 2014 | MAD14-68 | -14.7528 | 49.4875° |
| *Neohydrocoptus* sp. | SLE1545 | Kenya | Narok Co. | 2013 | SMB150613-B | -1.1533° | 35.2775° |
| *Liocanthodrus bicolor* | SLE1547 | Venezuela | Guárico | 2009 | VZ09-0110-02A | 8.1038° | -66.4371° |
| *Suphisellus puncticollis* | SLE1549 | USA | Florida | 2016 | SMB050816-A | 30.4739° | -85.9887° |
| *N. nanulus* * | SLE1127 | USA | Florida | 2016 | SMB060816-A | 30.0402° | 84.9836° |
| *N. sp. 6* * | SLE1649 | Suriname | Sipiliwini | 2016 | SR16-0316-01B | 4.6739° | -56.1847° |
| *N. gracilipes* ** | SLE2063 | Guatemala | Retalhuleu | 1974 | N/A | N/A | N/A |
| *N. gracilipes* | SLE1273 | Suriname | Brokopondo | 2017 | SR17-0322-02A | 4.9489° | -55.1804° |
| *N. gracilipes* | SLE2017 | Brazil | Amazonas | 2018 | BR18-0710-02B | -10.9176° | -62.3770° |
| *N. sp. 3* | SLE895 | Suriname | Sipiliwini | 2010 | SR10-0819-01A | 2.1753° | -56.7874° |
| *N. petrareptans* | SLE2055 | Suriname | Sipiliwini | 2010 | SR10-0901-01A | 2.1829° | -56.7873° |
| *N. petrareptans* | SLE2059 | Suriname | Sipiliwini | 2010 | SR10-0901-01A | 2.1829° | -56.7873° |
| *N. c.f.gracilipes* | SLE1635 | Brazil | Minas Gerais | 2018 | BR18-0228-02A | -15.1707° | -42.8035° |
| *N. c.f.gracilipes* | SLE1959 | Brazil | Amazonas | 2018 | BR18-0706-04A | -8.4410° | -61.6603° |
| *N. c.f.gracilipes* | SLE2074 | Brazil | Amapá | 2018 | BR18-0705-03E | -8.4376° | -61.7506° |
| *N. c.f.gracilipes* | SLE2049 | Suriname | Sipiliwini | 2013 | SR13-0824-03A | 3.7913° | -56.1495° |
| *N. traili* | SLE2028 | Brazil | Amapá | 2018 | BR18-0724-01A | 0.8090° | -51.9797° |
| *N. traili* | SLE2034 | Brazil | Amapá | 2018 | BR18-0722-03A | 1.9314° | -50.8610° |
| *N. traili* | SLE2020 | Brazil | Amapá | 2018 | BR18-0720-04A | 3.8116° | -51.7837° |
| *N. traili* | SLE1873 | Suriname | Sipiliwini | 2017 | SR17-0330-04A | 2.0040° | -55.9710° |
| *N. c.f.gracilipes* | SLE1879 | Suriname | Sipiliwini | 2017 | SR17-0401-01A | 2.0109° | -55.9845° |
| *N. traili* | SLE1885 | Brazil | Para | 2018 | BR18-0203-01E | -1.4929° | -54.5157° |
| *N. traili* | SLE1888 | Brazil | Para | 2018 | BR18-0203-01H | -1.4929° | -54.5157° |
| *N. traili* | SLE1639 | Brazil | Goiás | 2018 | BR18-0222-01A | -14.2774° | -47.7508° |
| *N. traili* | SLE1882 | Brazil | Mato Grosso do Sul | 2018 | BR18-0625-02A | -19.3062° | -57.6038° |
| *N. traili* | SLE2078 | Brazil | Mato Grosso do Sul | 2018 | BR18-0625-01A | -18.9518° | -57.6664° |
| *N. sabrouxi* | SLE2053 | Guadeloupe | Basse-Terre | 2013 | MM-190813-2 | 16.1799° | -61.6729° |
| *N. sabrouxi* | SLE909 | Guadeloupe | Basse-Terre | 2013 | MM-190813-2 | 16.1799° | -61.6729° |
| *N. traili* | SLE907 | Venezuela | Aragua | 2009 | VZ09-0104-02B | 10.3937° | -67.7960° |
| *N. traili* | SLE1636 | Venezuela | Bolívar | 2010 | VZ10-0713-03A | 7.3848° | -61.3253° |
| *N. traili* | SLE1534 | Brazil | Roraima | 2018 | BR18-0111-03B | 2.3837° | -60.5227° |
| *N. traili* | SLE1537 | Brazil | Roraima | 2018 | BR18-0111-03A | 2.3837° | -60.5227° |
| *N. traili* | SLE1876 | Brazil | Mato Grosso do Sul | 2018 | BR18-0622-03A | -20.4509° | -55.6218° |
| *N. c.f.gracilipes* | SLE2024 | Brazil | Rondonia | 2018 | BR18-0708-01A | -8.9081° | -62.1792° |
| *N. traili* | SLE1961 | Brazil | Bahia | 2018 | BR18-0224-03A | -11.6749° | -40.9964° |
| *N. traili* | SLE1641 | Brazil | Roraima | 2018 | BR18-0117-04A | 0.9131° | -59.5733° |
| *N. traili* | SLE1870 | Suriname | Sipiliwini | 2017 | SR17-0402-02B | 2.0040° | -55.9710° |
| *N. traili* | SLE1891 | Suriname | Sipiliwini | 2019 | SR19-0313-01B | 4.4060° | -57.2207° |
| *N. traili* * | SLE2065 | Brazil | Para | 1986 | Xingu River | -3.650° | -52.3667° |
| *N. traili* | SLE2057 | Guyana | Region 9 | 2013 | GY13-1031-01A | 2.1818° | -59.3384° |
| *N. traili* | SLE1631 | Guyana | Region 8 | 2014 | GY14-0319-02A | 5.3044° | -59.8376° |
| *N. traili* | SLE1644 | Suriname | Sipiliwini | 2010 | SR10-0904-01A | 2.36293° | -56.6977° |
| *N. traili* | SLE1632 | Suriname | Sipiliwini | 2012 | SR12-0310-02A | 2.4770° | -55.6290° |
| *N. traili* | SLE2061 | Suriname | Sipiliwini | 2010 | SR10-0819-01A | 2.1754° | -56.7874° |
| *N. traili* | SLE1633 | Suriname | Sipiliwini | 2013 | SR13-0813-04A | 3.7913° | -56.1494° |
| *N. traili* | SLE2036 | Suriname | Sipiliwini | 2010 | SR10-0819-01A | 2.1753° | -56.7874° |
| *N. traili* | SLE1642 | Guyana | Region 6 | 2014 | GY14-0925-01D | 4.1540° | -58.1771° |
| *N. traili* | SLE1638 | Suriname | Sipiliwini | 2012 | SR12-0727-03A | 4.7080° | -56.2193° |
| *N. traili* | SLE1640 | Guyana | Region 6 | 2104 | GY14-0928-01A | 4.1540° | -58.1771° |

**Table 4.**
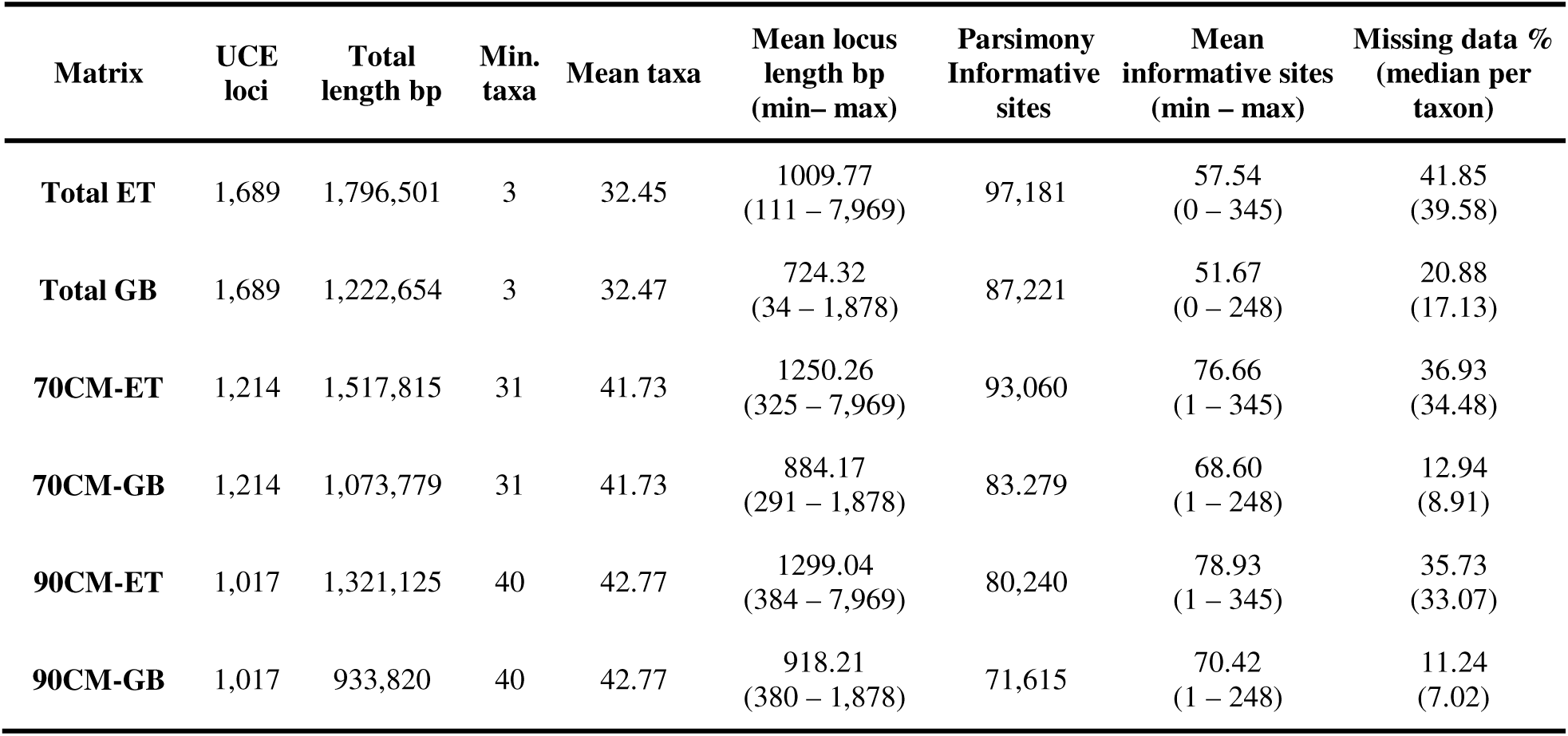
UCE alignment summary statistics. Alignment metrics for different trimming regimes and completeness thresholds for UCE alignments used in phylogenetic reconstructions. Left to right: Matrix: specifies output matrix from trimming method and completeness threshold treatments (CM=percent complete matrix, ET=edge-trimmed, GB=internal trimmed with gblocks); UCE loci: number of UCE loci represented in given alignment; Total length: total concatenated length of the alignment (bp); Min. taxa: minimum number of taxa required for a given UCE locus to be included in the respective alignment; Mean taxa: mean number of taxa represented in each locus for respective alignment; Mean locus length: mean length (in bp) of each locus in alignment; Informative sites: number of parsimony informative sites in alignments; Mean informative sites: mean number of informative sites per UCE locus; Missing data %: Mean percentage of missing data per taxon, combined undetermined (“?”) and gaps (“-”).

**Table 5.**
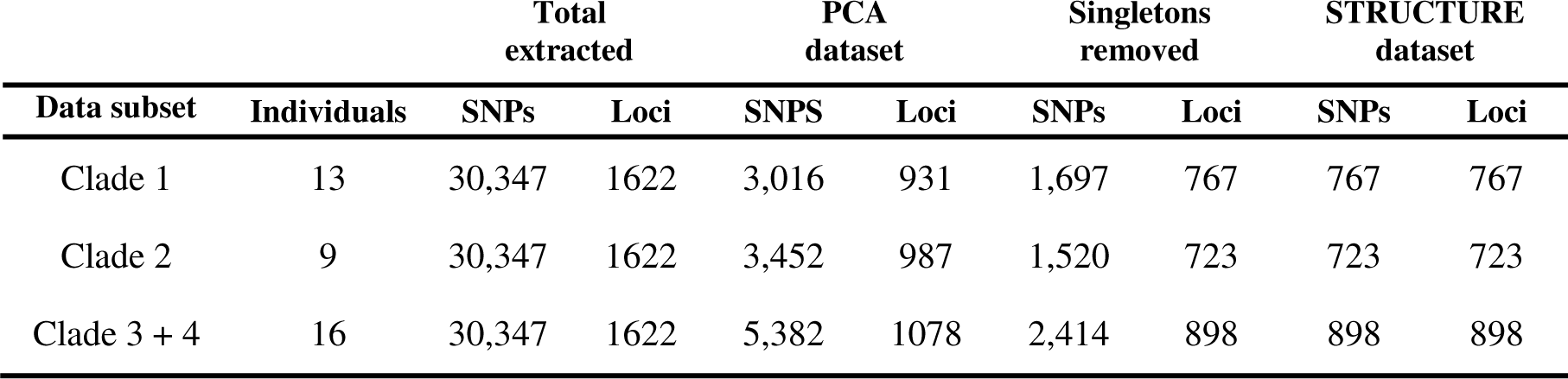
Summary statistics for SNP data subsets at different stages of SNP extraction and filtering from UCE reads. ‘Total extracted**’** indicates total number of SNPs and represented loci present in datasets following ‘PASS’ only QC filtering step and data subsetting (i.e. before processing in SNPfiltR); ‘PCA dataset’ indicates SNPs and loci present in datasets used for PCAs, prior to filtering for singletons and linkage; ‘Singletons removed’ indicates SNPs and loci present in datasets with a minimum minor allele count cutoff set to 2 to remove singleton data; ‘STRUCTURE dataset’ indicates unlinked SNPs (and loci) present in datasets used for STRUCTURE analyses.

Clade 2 includes several disjunct populations from south-central Brazil (Goiás, Mato Grosso do Sul), Guiana Shield and adjacent areas in Brazil and Venezuela (Roraima, Bolivar, Aragua), and Guadeloupe (Antilles) (Fig. 2, 3). These specific relationships do not conflict with Baca and Short (2020), however members of Clade 4 were there recovered as sister to Clade 2. It is again difficult to assess the effect of sparse sampling on patterns of clustering across these populations.

Clade 4 was recovered with two distinct sister populations in the Guiana shield with nearly complete overlap in distributions (Fig. 2c). The southernmost individual in these populations was from Para (SLE2065), a dried museum specimen collected near the type locality of *N. traili* (near Rio Xingu, Pará, Brazil). Samples of Clade 3, from western Amazonia and the Pantanal, and a single Clade 4 sample from the Cerrado were recovered as successive sisters to the more heavily sampled Guiana Shield populations. Clades 3 and 4 were not recovered by Baca and Short (2020), wherein these clades were intermixed, or individuals recovered elsewhere.

This is likely attributable to the greater amount of captured data with our UCE-based methods, as well as the resolution afforded by more complete taxon sampling (Heath et al., 2008), an effect seen in previous UCE investigations of Adephaga (Gustafson et al., 2019; Baca et al, 2021). The conflicting placement of SLE2024 between concatenated and ASTRAL analyses requires further investigation, with multiple potential causes.

### Phylogeography

The recovered topologies depict a repeated pattern in which northern South American populations have one or more successive sister populations in south central or western Brazil (Figs. 2, S2-S11). This signal is preserved across the different phylogenetic reconstruction methods utilized in this study (e.g ASTRAL and concatenated). Overall, the pattern is consistent with a process of diversification followed by dispersal into disjunct or widely dispersed populations, which may also potentially explain the divergence lineages of *N. gracilipes*. Evaluating the connectivity of these populations will require further investigation with sampling of the intermediate areas in the Amazon basin.

With Baca and Short (2020) estimating the crown age the whole *traili* group at ca. 13.2 mya and the age of the main complex (i.e. crown age of Clades 1+2+3+4) at ca. 7.7 ma, the evolution of this group would be subject to the dynamic environment of South American in the late Miocene through the present. During these times, however, the Guiana Shield and Brazilian Shield regions are thought to have provided relatively stable environments (Hoorn & Wesselingh, 2011 and works therein; Bicudo et al, 2019; Baker et al., 2020). It has been shown that communities in these areas comprise taxa that are phylogenetically over-dispersed (i.e., greater mean phylogenetic distance; Crouch et al., 2019; Bicudo et al., 2019). This is an indication that most diversification is not occurring *in situ* as is seen in more volatile areas such as lowland Amazonia (Bicudo et al., 2019), and communities are instead dispersal mediated (see Crouch et al., 2019). Here, we recover a pattern wherein many distinct lineages from disparate clades within the *traili group* inhabit the same regions, with the Guiana Shield having a notably high number of sympatric, yet genetically distinct populations. Note that this does not indicate that diversification is being primarily driven by dispersal into these areas; only that these areas may simply offer long-term habitat stability conducive to repeat invasions over longer-term evolutionary time. However, there are many potential geographic influences in the diversification of this group, for example climatic fluctuations in the Pleistocene, (Hoorn & Wesselingh, 2011; Baker et al., 2020 and citations therein; see also da Rocha & Kaefer, 2019). It is likely that multiple factors drive diversification in *traili group*. Parsing the influence of Neotropical geography beyond speculation requires a nuanced approach (Antonelli, 2018) that is beyond the current scope of this investigation.

### Taxonomic implications

The recovered patterns of phylogenetic and population genetic structure reveal that previous species delimitation hypotheses based on morphological traits and/or small numbers of loci (Young, 1978; Manuel, 2015; Baca and Short, 2018) are largely inconsistent with genome-wide patterns of relatedness throughout the distribution of this group. The evolution of patterns of elytral punctation provides a prime example of this discordance. Historically, *Notomicrus gracilipes* was diagnosed from *N. traili* and others by presenting punctation along the anteromedial margin of the elytra, along the suture (Fig. S2). As Fig. 2 depicts, specimens attributable to *N. gracilipes* (specimens labeled *N.* cf. *gracilipes*) are polyphyletic, showing that this character is a poor indicator of species boundaries. Thus, while *N. gracilipes* appears to be a valid species, sister to the rest of the *traili* group, the character formally used in identification is here shown unreliable. With other identifying characters such as the aedeagus or male protarsal claws showing subtle variation among clades, careful observation will be required to find correlating patterns of character state distributions across the traili phylogeny. Even then, parsing between species and populations morphologically may be difficult. Despite this cryptic morphology, the four currently valid species do not appear to require synonymy. Instead, it appears that additional species require description or in other cases, current species need to be expanded to include other populations. Properly attributing the name *N. traili* Sharp, 1882 to a clade will require careful investigation. The species was described from a single female type specimen (Sharp, 1882; Nilsson, 2011), which is problematic as females of *Notomicrus* lack the more easily identifiable primary sexual features of the aedeagus and secondary sexual modifications of the protarsal claws presented by males that have proven useful in *Notomicrus* taxonomy (e.g. Manuel, 2015; Guimarães and Ferreira-Jr, 2019). Observations of the female genitalia (Manuel, 2015; personal observations) show potential for characterizing lineages at the genus level in *Notomicrus*, but these characters appear to be too conserved to aid reliably in species delimitation. A further complication will be whether to resurrect *N. grouvellei* Régimbart, 1895 from synonymy with *N. traili*. This species was described from “Brésil: Mato Grosso” (Régimbart, 1895:18), an already large Brazilian state, which at that time included Mato Grosso do Sul.

A tentative undescribed species *N.* sp. 3 is clearly distinct from other members of the complex. To date it is known only from a few samples, but presents a distinct aedeagal morphology making readily identifiable with dissection. Externally, it appears similar to *N. gracilipes* or other punctate members.

## Conclusion

We successfully reconstructed the *N. traili* group phylogeny at the species and population level, supporting the efficacy of a tailored UCE approach for shallow-scale investigation and SNP-based population genetic analysis. Our reconstructions recovered multiple nested clades within the complex, each presenting a similar pattern of repeated diversification and dispersal. The results indicate that the Guiana Shield and Brazilian Shield appear to play an important role in providing stable environments for the persistence of *traili* lineages. Importantly, we provide a key foundation for future holistic investigations into the biogeographic factors of the diversification of the *traili* group and modified methodological framework for investigation at shallow evolutionary scales using a UCE-based approach. Finally, we find that that current classification does not appropriately represent the diversity of the *traili group*, which by all appearances contains more species than currently described. Further investigation will be needed to make taxonomic classification congruent with our newly bolstered understanding of the evolutionary relatedness within this group. As we continue to build on the robust dataset and methodological improvements presented herein, we are certain to gain more detailed understanding of *Notomicrus* diversity and South American evolutionary dynamics moving forward.

## Author Contributions

**Stephen Baca:** Conceptualization (co-lead); Funding acquisition (supporting); data capture and processing (lead); formal analyses (lead); methodology (lead); writing – original draft (lead); writing – review and editing (lead). **Grey Gustafson:** Conceptualization (supporting); methodology (supporting); writing – review and editing (supporting). **Devon DeRaad**: Methodology (supporting); writing – review and editing (supporting). **Alana Alexander**: Methodology (supporting); writing – review and editing (supporting). **Paul Hime:** conceptualization (supporting), methodology (supporting), writing – review and editing (supporting). **Andrew Short:** Conceptualization (co-lead); Funding acquisition (lead); writing – review and editing (supporting).

## Acknowledgements

We extend a heartfelt thanks to the Moyle Lab (University of Kansas) including Rob Moyle, Lucas Decicco and Emily Ostrow, for hosting the first author, and their generous gifts of time and insight on population genetics approaches used here. We additionally thank Rich Glor for advice on whole genome sequencing strategies for probe set design. We thank Michael Branstetter for insight on probe design approaches and help in the field. Huge thanks to KU Natural History Museum Biodiversity Institute molecular lab for facilitating all data capture, Riley Epperson and others at the KU Center for Research Computing for help and access with the High Performance Cluster, the KU-BI and affiliated PIs for generous access to computational nodes and storage, and members of the KU-BIMOL discussion group for their continued insight and shared learning experiences. Certainly not least, we thank our colleagues Michael Manuel for specimens, Mauricio Garcia (Venezuela), Luis Joly (Venezuela), Neusa Hamada (Brazil), Paul Ouboter (Suriname), Vanessa Kadosoe (Suriname), and WWF-Guianas for their facilitation of field sampling and the countries of Brazil, Guyana, Suriname and Venezuela for facilitating field sampling. Photo credits for images of *Neohydrocoptus* and *Sternocanthus* in Fig. 1: U. Schmidt (CC BY-SA 2.0; https://creativecommons.org/licenses/by-sa/2.0) accessed from https://www.flickr.com/search/?group_id=806927%40N20&view_all=1&text=Noteridae.

## Data availability

Sequence data captured for this project are deposited in the NCBI Sequence Read Archive under BioProject accession numbers provided in Table S2.

## Supplementary tables

**Table S1.** Proportions of missing data and gaps in concatenated alignments by sample. Values indicate missing data (‘?’), gaps (‘-’), and combined missing data and gaps (‘?’ + ‘-’) as percent of total respective alignment length. Samples are ordered by extraction code. Asterisks (*) indicate sample was dried museum specimen.

| Taxon | Sample | 70% CM-ET |  |  | 70% CM-GB |  |  | 90% CM-ET |  |  | 90% CM-GB |  |  |
| --- | --- | --- | --- | --- | --- | --- | --- | --- | --- | --- | --- | --- | --- |
|  |  | '?' | '-' | '?' + '-' | '?' | '-' | '?' + '-' | '?' | '-' | '?' + '-' | '?' | '-' | '?' + '-' |
| *N. sp. 3* | SLE895 | 9.48 | 25.66 | 35.14 | 9.07 | 2.28 | 11.36 | 8.58 | 25.80 | 34.38 | 7.96 | 2.27 | 10.23 |
| *N. traili* | SLE907 | 8.58 | 25.35 | 33.93 | 8.04 | 0.57 | 8.61 | 6.63 | 25.73 | 32.36 | 5.79 | 0.57 | 6.36 |
| *N. sabrouxi* | SLE909 | 12.38 | 24.47 | 36.85 | 11.85 | 0.54 | 12.39 | 9.54 | 25.31 | 34.85 | 9.03 | 0.55 | 9.58 |
| *N. nanulus* | SLE1127 | 46.63 | 16.67 | 63.31 | 47.77 | 6.74 | 54.51 | 44.16 | 17.34 | 61.50 | 45.23 | 7.06 | 52.29 |
| *N. gracilipes* | SLE1273 | 14.09 | 22.89 | 36.98 | 13.84 | 2.19 | 16.03 | 12.36 | 23.30 | 35.66 | 11.95 | 2.22 | 14.18 |
| *N. traili* | SLE1534 | 5.83 | 26.70 | 32.54 | 5.75 | 0.55 | 6.29 | 4.26 | 27.14 | 31.40 | 4.14 | 0.53 | 4.67 |
| *N. traili* | SLE1537 | 4.24 | 26.99 | 31.22 | 4.03 | 0.60 | 4.62 | 2.93 | 27.49 | 30.42 | 2.88 | 0.60 | 3.48 |
| *N. traili* | SLE1631 | 6.99 | 25.98 | 32.97 | 6.52 | 0.46 | 6.98 | 5.62 | 26.37 | 31.99 | 5.09 | 0.46 | 5.55 |
| *N. traili* | SLE1632 | 5.38 | 26.90 | 32.29 | 5.68 | 0.52 | 6.19 | 3.97 | 27.17 | 31.14 | 4.03 | 0.53 | 4.56 |
| *N. traili* | SLE1633 | 6.58 | 26.35 | 32.93 | 6.50 | 0.46 | 6.96 | 4.90 | 26.84 | 31.73 | 4.81 | 0.45 | 5.26 |
| *N. c.f.gracilipes* | SLE1635 | 7.66 | 25.83 | 33.50 | 7.12 | 0.73 | 7.85 | 6.42 | 26.10 | 32.52 | 5.70 | 0.70 | 6.40 |
| *N. traili* | SLE1636 | 9.69 | 24.93 | 34.62 | 8.59 | 0.54 | 9.13 | 6.64 | 25.92 | 32.56 | 5.70 | 0.53 | 6.23 |
| *N. traili* | SLE1638 | 5.87 | 26.90 | 32.76 | 6.10 | 0.46 | 6.56 | 4.02 | 27.28 | 31.30 | 4.02 | 0.45 | 4.48 |
| *N. traili* | SLE1639 | 7.53 | 26.21 | 33.74 | 7.37 | 0.62 | 7.99 | 5.64 | 26.82 | 32.46 | 5.58 | 0.60 | 6.18 |
| *N. traili* | SLE1640 | 6.04 | 26.03 | 32.07 | 5.39 | 0.48 | 5.87 | 4.89 | 26.18 | 31.07 | 3.99 | 0.48 | 4.48 |
| *N. traili* | SLE1641 | 5.40 | 26.24 | 31.64 | 5.05 | 0.57 | 5.62 | 3.98 | 26.78 | 30.76 | 3.76 | 0.54 | 4.30 |
| *N. traili* | SLE1642 | 11.17 | 24.27 | 35.44 | 10.21 | 0.53 | 10.74 | 9.02 | 24.96 | 33.98 | 8.15 | 0.55 | 8.70 |
| *N. traili* | SLE1644 | 6.97 | 25.88 | 32.85 | 6.36 | 0.49 | 6.85 | 4.77 | 26.82 | 31.59 | 4.56 | 0.48 | 5.03 |
| *N. c.f.gracilipes* | SLE1649 | 20.24 | 20.15 | 40.40 | 20.03 | 2.69 | 22.72 | 18.45 | 20.61 | 39.06 | 18.29 | 2.70 | 20.99 |
| *N. traili* | SLE1870 | 7.71 | 25.64 | 33.36 | 7.05 | 0.65 | 7.70 | 6.22 | 26.11 | 32.32 | 5.53 | 0.64 | 6.17 |
| *N. traili* | SLE1873 | 10.73 | 25.35 | 36.08 | 10.61 | 0.70 | 11.30 | 8.86 | 25.68 | 34.54 | 8.46 | 0.69 | 9.14 |
| *N. traili* | SLE1876 | 10.30 | 25.29 | 35.59 | 9.88 | 0.77 | 10.65 | 8.69 | 25.88 | 34.57 | 8.42 | 0.76 | 9.18 |
| *N. c.f.gracilipes* | SLE1879 | 10.58 | 25.14 | 35.72 | 9.95 | 0.71 | 10.66 | 8.94 | 25.52 | 34.46 | 8.15 | 0.70 | 8.86 |
| *N. traili* | SLE1882 | 8.04 | 26.15 | 34.20 | 7.97 | 0.66 | 8.63 | 6.60 | 26.49 | 33.09 | 6.34 | 0.66 | 6.99 |
| *N. traili* | SLE1885 | 8.68 | 25.80 | 34.48 | 8.19 | 0.73 | 8.91 | 7.21 | 26.19 | 33.40 | 6.67 | 0.70 | 7.37 |
| *N. traili* | SLE1888 | 13.97 | 24.22 | 38.19 | 13.41 | 0.69 | 14.11 | 12.88 | 24.40 | 37.27 | 12.07 | 0.69 | 12.76 |
| *N. traili* | SLE1891 | 12.03 | 23.96 | 35.99 | 10.64 | 0.61 | 11.24 | 10.82 | 24.54 | 35.36 | 9.71 | 0.61 | 10.33 |
| *N. traili* | SLE1959 | 10.86 | 25.16 | 36.02 | 10.44 | 0.73 | 11.17 | 9.00 | 25.75 | 34.76 | 8.62 | 0.74 | 9.36 |
| *N. traili* | SLE1961 | 8.15 | 25.91 | 34.06 | 8.05 | 0.55 | 8.60 | 5.65 | 26.71 | 32.36 | 5.64 | 0.55 | 6.19 |
| *N. gracilipes* | SLE2017 | 15.21 | 22.45 | 37.67 | 14.45 | 2.21 | 16.66 | 12.65 | 23.17 | 35.82 | 11.90 | 2.28 | 14.18 |
| *N. traili* | SLE2020 | 8.33 | 26.31 | 34.64 | 8.56 | 0.74 | 9.31 | 7.38 | 26.45 | 33.82 | 7.39 | 0.74 | 8.14 |
| *N. c.f.gracilipes* | SLE2024 | 19.78 | 21.19 | 40.97 | 17.29 | 0.84 | 18.13 | 19.51 | 21.44 | 40.95 | 17.23 | 0.82 | 18.06 |
| *N. traili* | SLE2028 | 7.88 | 26.07 | 33.95 | 7.71 | 0.74 | 8.44 | 6.46 | 26.32 | 32.78 | 6.07 | 0.73 | 6.80 |
| *N. traili* | SLE2034 | 5.61 | 26.66 | 32.27 | 5.24 | 0.71 | 5.95 | 4.37 | 26.85 | 31.21 | 3.74 | 0.69 | 4.43 |
| *N. traili* | SLE2036 | 8.13 | 25.91 | 34.04 | 7.92 | 0.48 | 8.40 | 6.80 | 26.27 | 33.07 | 6.56 | 0.46 | 7.02 |
| *N. c.f.gracilipes* | SLE2049 | 5.99 | 26.65 | 32.64 | 5.81 | 0.73 | 6.55 | 4.85 | 26.97 | 31.81 | 4.64 | 0.72 | 5.36 |
| *N. sabrouxi* | SLE2053 | 6.72 | 26.16 | 32.88 | 6.47 | 0.59 | 7.06 | 4.95 | 26.75 | 31.70 | 4.84 | 0.60 | 5.44 |
| *N. petrareptans* | SLE2055 | 7.93 | 25.95 | 33.88 | 7.61 | 0.78 | 8.39 | 5.79 | 26.77 | 32.57 | 5.64 | 0.78 | 6.43 |
| *N. traili* | SLE2057 | 9.35 | 25.34 | 34.69 | 8.80 | 0.46 | 9.26 | 8.21 | 25.63 | 33.84 | 7.52 | 0.46 | 7.97 |
| *N. petrareptans* | SLE2059 | 8.57 | 26.25 | 34.82 | 8.83 | 0.78 | 9.61 | 6.30 | 26.71 | 33.01 | 6.27 | 0.79 | 7.06 |
| *N. traili* | SLE2061 | 7.22 | 25.98 | 33.20 | 6.84 | 0.45 | 7.29 | 5.18 | 26.71 | 31.89 | 4.97 | 0.43 | 5.41 |
| *N. gracilipes\** | SLE2063 | 74.36 | 6.96 | 81.32 | 72.16 | 2.35 | 74.52 | 73.16 | 7.44 | 80.60 | 70.99 | 2.54 | 73.53 |
| *N. traili\** | SLE2065 | 52.00 | 11.19 | 63.19 | 48.35 | 0.55 | 48.89 | 50.87 | 11.58 | 62.46 | 47.31 | 0.55 | 47.86 |
| *N. c.f.gracilipes* | SLE2074 | 9.93 | 25.74 | 35.67 | 10.04 | 0.72 | 10.76 | 8.29 | 26.03 | 34.32 | 8.09 | 0.74 | 8.83 |
| *N. traili* | SLE2078 | 12.10 | 25.06 | 37.16 | 12.23 | 0.57 | 12.80 | 9.26 | 26.00 | 35.26 | 9.52 | 0.61 | 10.13 |
| **Mean** |  | 12.47 | 24.46 | 36.93 | 11.99 | 0.94 | 12.94 | 10.79 | 24.94 | 35.73 | 10.29 | 0.95 | 11.24 |
| **Median** |  | 8.57 | 25.80 | 34.48 | 8.19 | 0.65 | 8.91 | 6.64 | 26.11 | 33.07 | 6.34 | 0.64 | 7.02 |

**Table S2.** NCBI sequence read archive (SRA) accession numbers for UCE raw sequence reads.

| **Taxon** | **Sample code** | **SRA accession** |
| --- | --- | --- |
| *Neohydrocoptus sp.* | SLE1545 | XX |
| *Liocanthyrus bicolor* | SLE1547 | XX |
| *Sternocanthus sp.* | SLE1544 | XX |
| *Suphisellus puncticollis* | SLE1549 | XX |
| *N. nanulus* | SLE1127 | XX |
| *N. sp. 6* | SLE1649 | XX |
| *N. gracilipes* | SLE2063 | XX |
| *N. gracilipes* | SLE1273 | XX |
| *N. gracilipes* | SLE2017 | XX |
| *N. sp. 3* | SLE895 | XX |
| *N. petrareptans* | SLE2055 | XX |
| *N. petrareptans* | SLE2059 | XX |
| *N. c.f.gracilipes* | SLE1635 | XX |
| *N. c.f.gracilipes* | SLE1959 | XX |
| *N. c.f.gracilipes* | SLE2074 | XX |
| *N. c.f.gracilipes* | SLE2049 | XX |
| *N. traili* | SLE2028 | XX |
| *N. traili* | SLE2034 | XX |
| *N. traili* | SLE2020 | XX |
| *N. traili* | SLE1873 | XX |
| *N. c.f.gracilipes* | SLE1879 | XX |
| *N. traili* | SLE1885 | XX |
| *N. traili* | SLE1888 | XX |
| *N. traili* | SLE1639 | XX |
| *N. traili* | SLE1882 | XX |
| *N. traili* | SLE2078 | XX |
| *N. sabrouxi* | SLE2053 | XX |
| *N. sabrouxi* | SLE909 | XX |
| *N. traili* | SLE907 | XX |
| *N. traili* | SLE1636 | XX |
| *N. traili* | SLE1534 | XX |
| *N. traili* | SLE1537 | XX |
| *N. traili* | SLE1876 | XX |
| *N. c.f.gracilipes* | SLE2024 | XX |
| *N. traili* | SLE1961 | XX |
| *N. traili* | SLE1641 | XX |
| *N. traili* | SLE1870 | XX |
| *N. traili* | SLE1891 | XX |
| *N. traili* * | SLE2065 | XX |
| *N. traili* | SLE2057 | XX |
| *N. traili* | SLE1631 | XX |
| *N. traili* | SLE1644 | XX |
| *N. traili* | SLE1632 | XX |
| *N. traili* | SLE2061 | XX |
| *N. traili* | SLE1633 | XX |
| *N. traili* | SLE2036 | XX |
| *N. traili* | SLE1642 | XX |
| *N. traili* | SLE1638 | XX |
| *N. traili* | SLE1640 | XX |

